# Forensic STR profiling using Oxford Nanopore Technologies’ MinION sequencer

**DOI:** 10.1101/433151

**Authors:** Senne Cornelis, Sander Willems, Christophe Van Neste, Olivier Tytgat, Jana Weymaere, Ann-Sophie Vander Plaetsen, Dieter Deforce, Filip Van Nieuwerburgh

## Abstract

Forensic STR profiling using massively parallel sequencing (MPS) has gained much attention as an alternative for the traditional capillary electrophoresis (CE) methods. Oxford Nanopore Technologies recently developed the ‘MinION’, a pocket-sized nanopore sequencer with promising features that could be useful in the field of forensic genetics. We applied this technology for forensic sequencing in a pilot study. Using standard STR primers, originally designed for multiplex PCR and CE, we developed a library preparation method suited for nanopore sequencing. Several analysis approaches were evaluated to explore the usefulness of the generated data: we developed and applied both a sequence-based and an amplicon length-based analysis on data originating from a 14-loci multiplex PCR on a single contributor DNA sample. Despite the high sequencing error rate, the analyses yielded partial forensic profiles with some useful evidential value.

## 1 Introduction

Short tandem repeat (STR) profiling using multiplex PCR and subsequent capillary electrophoresis (CE) is still the state-of-the-art and most frequently used method to perform DNA based human identification [^1^]. However, in recent years several sequencing based approaches for forensic human identification have been developed and are being commercialized as streamlined, validated methods [^2–4^]. The use of sequencing for analysis of forensic amplicons has several well described advantages over CE [^5–8^]. A virtually unlimited number of autosomal, X-, Y-STR and SNP loci can be multiplexed in a single experiment, only being limited by the PCR multiplex capability. Alleles with an identical length but a different sequence can be distinguished, leading to more discriminative results. Despite these advantages, forensic sequencing based approaches are not yet widely used. The purchase of a state-of-the-art sequencer is a substantial investment. Moreover, the cost per sample is much higher compared to conventional CE analysis, even when a substantial number of samples are pooled during sequencing. Nanopore sequencing, an alternative sequencing technology, holds some promise to become the technology of choice to perform forensic STR profiling [^9^]. Oxford Nanopore Technologies (ONT) commercialized the pocket-sized, USB-powered MinION nanopore sequencer. While the cost of the MinION sequencer is merely a $1000, the cost of the disposable flow cells is currently prohibitive for forensic sequencing. However, this cost is predicted to become feasible in the coming years. The VolTRAX add-on is a rapid, programmable, portable, disposable sample processor, which is designed to convert a biological sample to a library that is compatible with nanopore analysis. The entire workflow is performed without any need for human intervention. ONT’s propitiatory analysis software allows performing on-the-fly analysis with the computing power of a smartphone and an internet connection. This makes it possible to sequence biological samples virtually anywhere, also in places where access to lab facilities is limited such as developing countries and regions hit by disasters such as an Ebola outbreak [^10^]. The possibility to work independently from a laboratory would also make it feasible to perform human identification by sequencing at border crossings, police check-points, police stations etc. without the need for highly qualified personnel. In order to allow a DNA sample to be ONT sequenced, a library preparation is required to ligate a ‘leader’ sequence to one end of the DNA fragments and a specific hairpin sequence to the other end. Theoretically, no PCR is required as individual DNA molecules are sequenced when they pass through the nanopores. However, a multiplex PCR is needed to amplify the forensic loci to an amount that is suitable for the library preparation (1 *µ*g is recommended as input). At the time of our study, only DNA fragments longer than 100 base pair (bp) could be analyzed. As some of the current STR amplicons are close to or shorter than this minimum length, we circumvented this problem by concatenating the PCR amplicons using a ligation step prior to the library preparation. During data-analysis, the original amplicon reads were retrieved as subreads from the read produced by the MinION sequencer, using a custom software pipeline. Several data-analysis approaches were evaluated to characterize the usefulness of the data originating from a 14- loci multiplex PCR on a single contributor DNA sample. Based on exploratory results generated with previously developed forensic tools such as MyFLq (My-Forensic-Loci-Queries) [^5,7^], both a sequence-and an amplicon length based approach were developed. In the length-based approach, we generated a CE-like length-based profile, using the length of the retrieved subreads. In the sequence-based method, amplicon sequences were aligned against a reference database containing all known al-lele sequences, with use of lenient nanopore-data-specific parameters. Particular attention was paid to the sex marker amelogenin. The X and Y alleles differ by seven SNVs (single nucleotide variants) and a six bp indel. A comparison of these alleles enables us to assess the ability of the MinION to identify SNVs to discriminate forensic alleles.

## 2 Materials & Methods

### 2.1 Nanopore library preparation and sequencing

The results presented in this paper were obtained from sample 9948A, a male single contributor control DNA sample (Promega, Madison, US). The reference profile for the 9948A sample is shown in Supplementary Table 1. The sample’s STR loci were amplified in a 14-plex PCR (primer sequences available in Supplementary Table 2) using the HotStarTaq Master Mix Kit (Qiagen). Initial denaturation at 95°C for 15 min was followed by 30 amplification cycles consisting of denaturation at 95°C for 60 s, primer annealing at 59°C for 60 s and elongation at 72°C for 80 s. Subsequent to this amplification a final elongation step of 10 min at 72°C was performed. The resulting PCR amplicons were end-polished using the NEBNext End-Repair module (NEB, Ipswich, USA) and cleaned up using 1X volume AMPure^®^ XP beads (Beckman Coulter, High Wycombe, UK). Random concatenation of the generated amplicons was performed using the Blunt T/A Ligase Mastermix (M0367S NEB, Ipswich, USA). The ligation reaction proceeded for 15 min, after which a cleanup with AMPure XP beads was performed. The ligation efficiency was verified by gel-electrophoresis (E-gel 2 % Thermo Fisher) and 12K Agilent Bioanalyzer chip (Agilent Technologies, Santa Clara, US). Prior to the library preparation, the ligated PCR amplicons were quantified using a Qubit fluorometer (Life Technologies, Paisley, UK) and diluted to a final concentration of 45 ng/*µ*l. The Oxford Nanopore library preparation protocol requires the attachment of a leader and hairpin (HP) adaptor to the sample. End-repair was performed on 1.980 *µ*g of the previously ligated PCR amplicons using the Ultra End-Repair/dA-Tailing module (NEB, Ipswich, USA) according to the manufacturer’s instructions. The resulting A-tailed DNA was cleaned up using 1.8X volume AMPure XP beads (Beckman Coulter, High Wycombe, UK) according to the manufacturer’s instructions. 50 *µ*l Blunt/TA ligase Mastermix (NEB) was added to the eluate, followed by 8 *µ*l nuclease free water, 10 *µ*l Adaptor Mix (AMX; including the leader sequence) and 2 *µ*l HP adaptor. The library was mixed by inversion between each sequential addition. After incubating for 10 minutes at room temperature 1 *µ*l of HP tether was added. Subsequently, the reaction was left to proceed for 10 min at room temperature. This adaptor-ligated, tether-bound library was purified using 500 ng MyOne C1 Dynabeads (Thermo Fisher, UK). The sample was resuspended in 25 *µ*l elution buffer and left to incubate for 10 min at room temperature, before pelleting the MyOne beads and collecting the supernatant containing the library. Finally, the library was quantified fluorimetrically as mentioned above. This so-called pre-sequencing mix had a final DNA concentration of 17.6 ng/*µ*l. 6 *µ*l of this pre-sequencing mix was supplemented with 4 *µ*l fuel mix and 75 *µ*l running buffer, after which the mixture was further diluted with 150 *µ*l water to yield the actual sequencing mix. For sequencing, a new flow cell (MkI, R7) was used to ensure optimal pore availability. The 48-hour MAP-006 sequencing protocol was initiated using the MinKNOW control software. The flow cell was topped up with newly made sequencing mix (as described above) after 6 h and the protocol was stopped after 12 h of non-stop sequencing.

### 2.2 Base calling and locus sequence extraction

To process the *fast5* files generated by the MinION sequencer, a custom open source python script had to be developed. These *fast5* files contain reads that originate from the randomly ligated STR amplicons. The reads need to be split into subreads representing individual STR amplicons, which must then be allocated to the correct individual loci. The analysis consisted of the following steps: (1) Sequences and associated sequencing quality scores were extracted from the *fast5* files and were written into a *fastq* file. (2) Using the primer sequences (Supplementary Table 2) the subreads within each read were identified and allocated to a specific locus if at least a unique 6 bp primer sequence fragment of the forward and reverse primers was recognized. These 6 bp fragments were considered unique, when they only occurred once in a specific primer, and were not present in any other primer or reference allele sequence. Additionally, the subreads were filtered for a minimum length (50bp) in order to retain only full-length amplicon sequences and exclude short artifacts. (3) The filtered subreads were merged into a single *fasta* file and the assigned locus was added to each subread’s description line. All downstream data analysis approaches are based on these subreads. Detailed documentation including the python script is available on the Nanopore Forensic (NanoFore) Github repository (https://github.com/SenneC1/NanoFore).

### 2.3 Sequence-based profiling

In the sequence-based profiling approach, the subreads were aligned against an allele database using the Burrows-Wheeler Aligner (BWA-MEM) algorithm and the ONT2D option, saving the subsequent results in a SAM file. The allele database was constructed using information from the Short Tandem Repeat DNA Internet Database (STRbase) [^11^] of the NIST Standard Reference Database. The BWA-MEM ONT2D option was specifically designed for mapping of nanopore reads, allowing a higher indel rate. To ensure only uniquely mapped subreads were used in downstream analysis, all alignments with a mapping score of 0 (not mapped uniquely) or a ‘flag’ of 4 (not mapped) were filtered from the SAM file. For each allele in the allele sequence database, the corresponding mapped subreads were counted. The relative frequencies per locus were calculated and plotted to create a CE-like profile. The amelogenin locus, a gender identification marker, is unlike the other loci investigated in this multiplex no STR locus. The amelogenin X and Y loci differ by seven SNVs and a six bp indel. In order to assess whether these SNVs and the indel could be detected reliably, 10 random subreads, which uniquely mapped against the AmelX sequence were forced to map against the AmelY sequence. The mapping result was visualized using IGV viewer [^12^]

### 2.4 Length-based profiling

In the second data-analysis approach, the length-based approach, a CE-like length-based profile was generated. The *fasta* output was used to calculate the length of each subread. For each locus, a histogram of the subread sequence length was generated using python’s matplotlib module [^13^]. In each of the length-based histograms, the two maxima were identified. In case the second maximum was less than 20 % of the main peak, it was not taken into consideration. No more than two maxima were allowed as the sample is known to be from a single contributor. For each of the detected maxima, a Gaussian curve was fitted. In case two peaks were detected, two overlapping Gaussian curves were fitted. A documentation of the procedure including the used python script can be found on the NanoFore Github repository (https://github.com/SenneC1/NanoFore). For each Gaussian fitting, the amplitude and standard deviation (SD) were calculated.

### 2.5 Evidential value of the profile

To assess the evidential value of the generated STR profiles, an RMNE value was calculated using the methods described by Van Nieuwerburgh *et al* [^14^], Van Neste *et al* [15] and Willems *etal* [^16^]. In the length-based profile, the detected peaks span several possible alleles. The true alleles and the zygosity cannot be exactly deduced from the profile, but results show that the true alleles lie within one standard deviation of the peak average. Without prior knowledge, it is impossible to know how many persons contributed to the profile. For the RMNE calculation, the alleles within the range of the peak average ± 1 standard deviation (SD), are considered as observed. The RMNE value is calculated as if the profile is a mixed profile with these observed alleles. The allele frequencies used in the calculation were the European allele frequencies obtained from the *popSTR* Europe database (spsmart.cesga.es/popstr.php). Allele frequencies of zero were assigned a value of 0.001 to avoid that the presence of these rare alleles in a profile would infer too much evidential value. A similar analysis was performed for the sequence-based profile: For each locus, the allele associated with the maximum number of mapped subreads as well as the alleles with a number of mapped subreads higher than 50% of the maximum, are considered as observed. Because not all true alleles are correctly detected, the RMNE value is calculated allowing for a number of drop-outs.

## 3 Results

### 3.1 Nanopore library preparation

The ligation protocol produced DNA fragments with a median length of around 2000 bp (Fig. 1 A). The actual nanopore library preparation yielded 440 ng of library, which is in accordance with ONT documentation. The sequencing proceeded for 12 h and produced 61003 sequence reads. Of these, 22712 reads were categorized as high quality two-directional (2D) reads. Fig. 1 B) shows the read length histogram of these high quality reads. The median read length was 1187 bp. The longest sequence measured was 9878 bp. Subreads (see Locus sequence extraction) were extracted from these high quality 2D reads yielding a *fasta* file with 16729 subreads that are allocated to their respective loci. Supplementary Table 3 shows the subread count per locus. In general, all loci are represented in the same order of magnitude indicating a uniform amplification and sequencing. One locus, namely SE33, is slightly under-represented.

**Figure 1:**
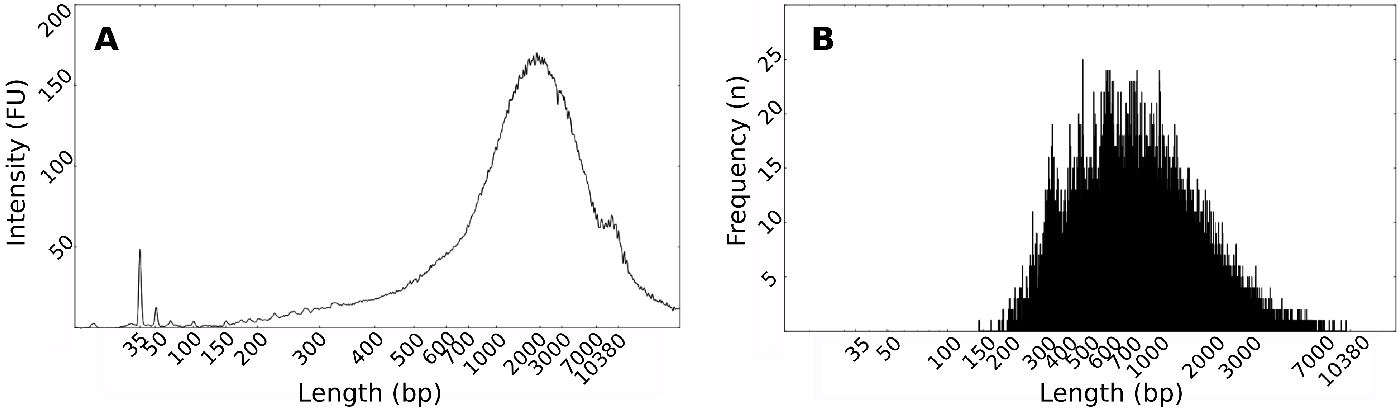
**A)** Length profile (bp)of concatenated amplicons as measured with 12K Agilent Bioanalyzer chip (logscale), internal marker at 35 bp and 10380 bp. **B)** Read length (bp) histogram of the high quality passed two-directional(2D) nanopore reads.

### 3.2 Sequence-based profiling

Of the 16729 subreads, 4222 could be uniquely mapped against the reference allele database. The relative frequency of the mapped subreads per locus is shown in Fig. 2. (The absolute frequencies are tabulated in Supplementary Table 4). Green colored bars indicate the true alleles. For most loci (D3S1358, D5S818, D7S820, D13S317, D18S51, D21S11, FGA, D16S539, TPOX, vWA) the majority of the subreads maps with the true alleles. However, for other loci (D8S1179, SE33, TH01), a relatively large number of the subreads map with several allele sequences which are not true alleles, thereby creating noise. Some heterozygous loci (D3S1358, D5S818, FGA, D18S51) show a good balance between both alleles while others (TPOX, D21S11) are skewed.

**Figure 2:**
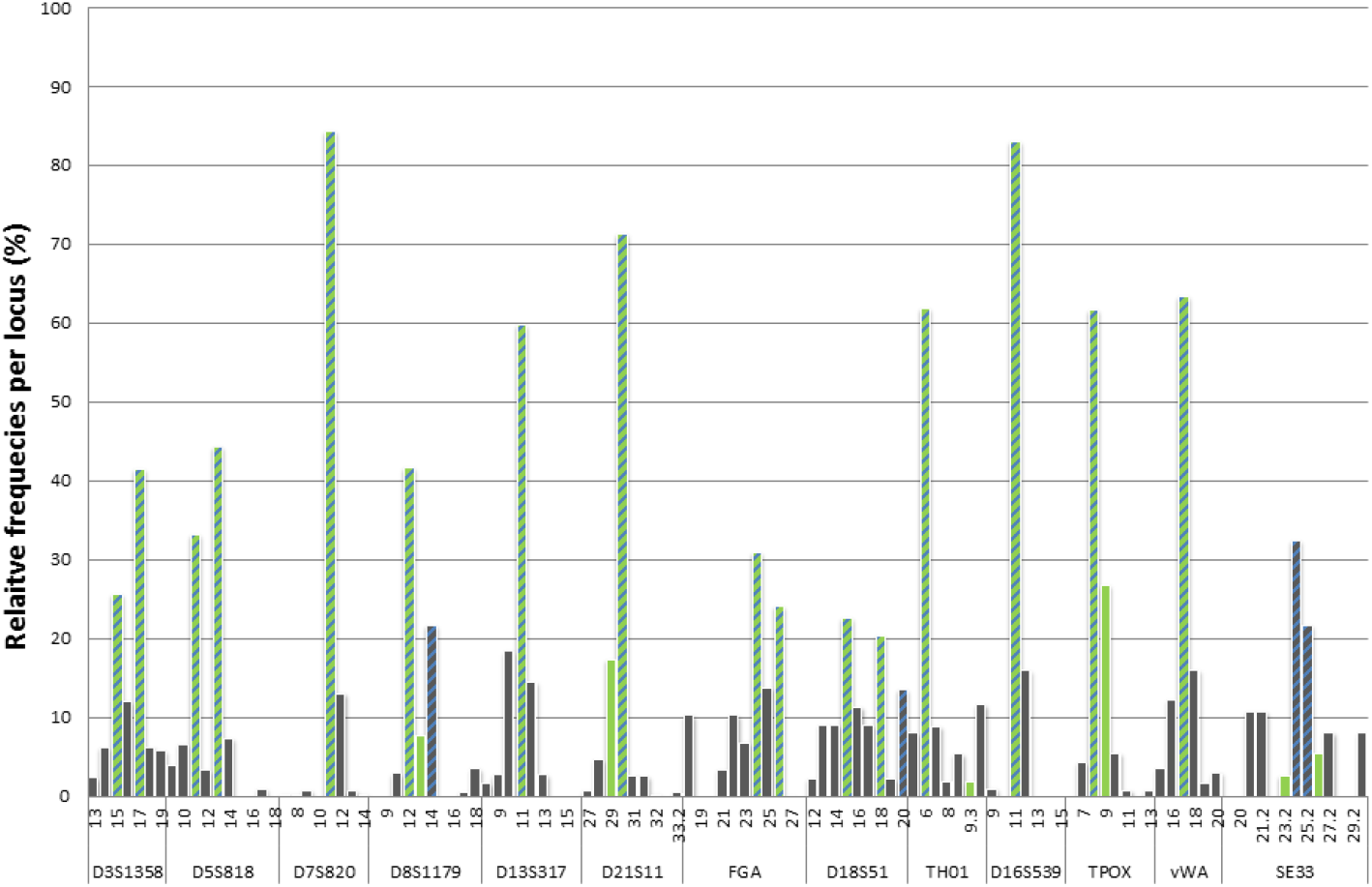
Relative frequency of uniquely mapped subreads per locus. All true alleles are indicated in Green. The blue patterning indicates when an allele is considered as observed. Hence, Green bars with blue patterning indicate correctly genotyped alleles. Green bars without patterning indicate a failure to identify the correct allele, a drop-out. Gray bars with blue patterning indicate an incorrect allele was observed, a drop-in

To call alleles as observed and generate an STR profile, the following rule was implemented: for each locus, the allele associated with the maximum number of mapped reads is considered as observed. Furthermore, additional alleles are considered as observed if the number of mapped reads is higher than 50% of the maximum. All alleles considered as observed based on this rule are indicated with a blue pattern in Fig. 4. Seven loci (D13S317, D3S1358, DS5818, D7S820, D16S539, FGA, vWA) were correctly genotyped, three loci had a drop-out (D21S11, TH01 and TPOX), one locus had a drop-in (D18S51), one locus had a wrongly called allele (D8S1179; counting for 1 drop-out and 1 drop-in) and for one locus two wrong alleles were called (SE33; counting for 2 dropouts and 2 drop-ins). The resulting profile is shown in Supplementary Table 1. Besides the 13 STR loci, the amelogenin locus was also included in the sequencing study. The two possible amelogenin alleles (AmelX and AmelY), which differ from each other in seven single nucleotide variants (SNVs) and a six bp indel, could be observed. Using the BWA–MEM ONT2D algorithm, 325 subreads mapped uniquely with the AmelX reference, whereas 373 subreads uniquely mapped the AmelY reference. This is close to the theoretically expected 1:1 ratio, for a male sample. Of the 779 subreads, 81 could not be matched uniquely to one of both reference sequences. The differences between the AmelY reference and the AmelX mapped subreads were visualized in Fig. 3. Here, 10 subreads, which uniquely mapped to the AmelX allele, were forced to map to the AmelY reference, thereby highlighting the mismatches. The most obvious difference is the large deletion situated between the 40 and 60 bp mark on Fig. 3. This deletion corresponds in both length and position to the known six bp indel. Moreover, seven positions show a high number of mismatch nucleotides. Again, this is in concordance with the position and the sequence of the seven SNVs that are known to be different between amelogenin X and Y alleles.

**Figure 3:**
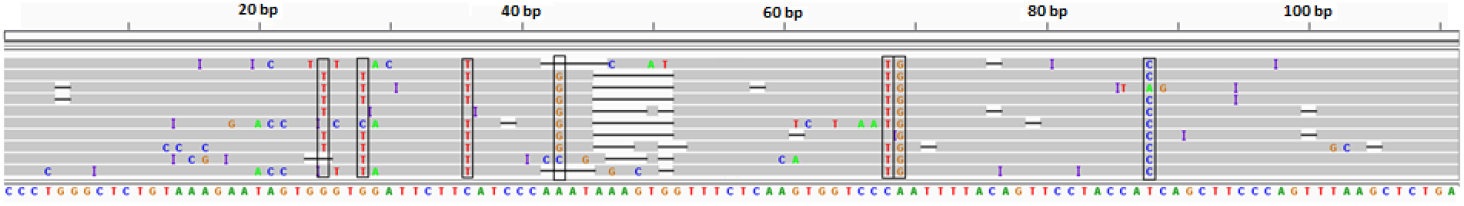
Amelogenin X subreads (above) mapped against the Amelogenin Y reference (below). Black boxes indicate the seven known single nucleotide variants between AmelX and AmelY. Mismatching nucleotides are indicated separately with their respective letter.

**Figure 4:**
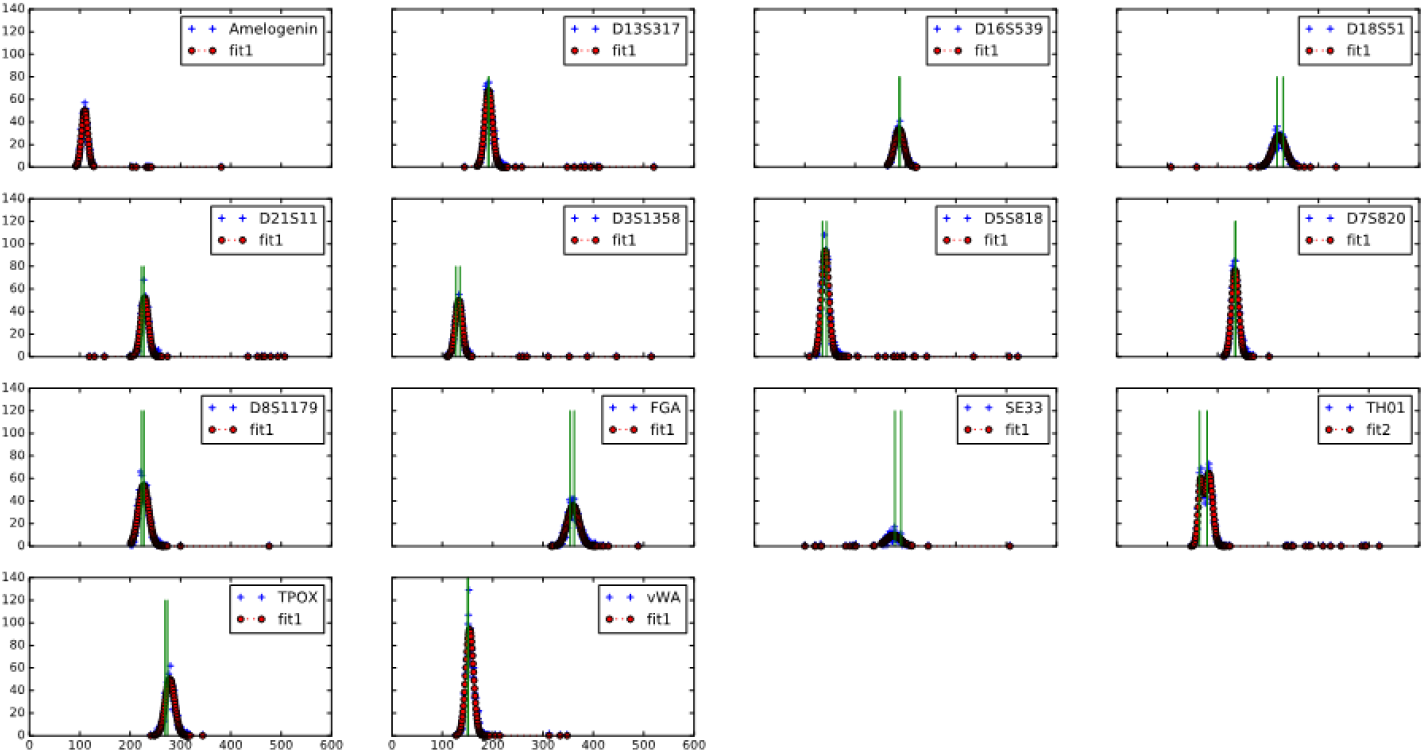
Gaussian fittings of the sequence length histograms of each locus (red). The length of the true alleles per locus is indicated green. All plots are scaled equally.

### 3.3. Length-based profiling

The subread length histograms of all analyzed loci are shown in Supplementary Table 5. In an attempt to obtain the correct allele determination for individual STR loci using sequence length data, Gaussian curves were fitted to these length distributions. The results of these Gaussian fittings are shown in Fig. 4. Of all heterozygous loci, only the TH01 locus was fitted with two overlapping Gaussian curves. The subread length corresponding to the mean ± 1 SD of the fitted Gaussian, and the corresponding range of observed alleles is shown in Supplementary Table. 5. The mean of the Gaussian function is a good predictor for the allele length of the homozygous loci. The true repeat length of D13S317 (11:11) and D7S820 (11:11) corresponds perfectly with the repeat length calculated using the mean of the Gaussian curve. For the homozygous loci D16S539 (11:11) and vWA (17:17), the calculated repeat length only differs one nucleotide (10.3) and three nucleotides (17.3) respectively. All true alleles of the different heterozygous loci are included within the mean ± 1 SD of the Gaussian function. The only exception is allele 26.2 of locus SE33 that is one bp outside of this range. The resulting profile is shown in Supplementary Table 1.

### 3.4. Evidential value of the profile

All observed alleles, from both the sequence -and length-based profiling methods are shown in Supplementary Fig. 1. Compared to the true profile, the observed sequence based profile has 4 drop-ins and 6 drop-outs. The resulting RMNE value, allowing for 6 drop-outs is 5.32E-7 or 1 in 1,880,124. The RMNE value of the length-based profile was calculated including all alleles within the mean ± 1 SD of the Gaussian function (see Methods section) and was 0.00026 or 1 in 3,846.

## 4 Discussion

The yield obtained after 12 consecutive hours of Nanopore sequencing is in line with those reported by other ONT community members, taking into account that only 184 pores were available for sequencing. The ligation-mediated concatenation of small STR amplicons into longer strands circumvents the minimum length requirement set by the base calling software. At the time of our study, the base calling software required at least 150 template and 150 complement events in order to perform a 2D workflow and generate a 2D read. Without concatenation, a substantial portion of the STR amplicons would not have been analyzed. The base calling software is constantly updated and improved: In the meanwhile, the minimum DNA fragment length needed to produce a read has decreased to 100 bp. Still, the concatenation approach is useful, as it ensures the possibility to analyze even shorter miniSTR and SNP amplicons.

In a first attempt to extract a forensic profile from the sequencing data, the open source MyFLq application was used [^5,7^] (data not shown). This program counts and stacks reads that have an identical sequence or identical length. The software did not provide satisfactory results, most likely because it was designed for low error rate sequences such as Illumina reads. To improve upon these exploratory results, both a sequence-and a length-based approach were developed to extract useful forensic results from the nanopore data.

### 4.1. Sequence-based profiling

The sequence-based profiling approach relies on the unique alignment of the sequencing reads against an allele database. However, a high percentage of subreads (63.1%-97.3%) maps with an equal score to more than one allele present in the database. Fortunately, these ambiguously mapped subreads can be filtered. The majority of the remaining uniquely mapped subreads align with the true alleles. Nevertheless, a substantial part of subreads aligns with an incorrect allele. These incorrect mappings can be attributed by two factors: 1) The high amount of sequencing errors produced by the MinION sequencer which results in a more challenging alignment. 2) The reference database consists of alleles from the same locus, which display a high level of similarity between them. Hence, the existing sequencing errors can easily result in identical mapping scores. This problem is further exacerbated with larger reference databases. The SE33 locus, which showcases the largest database, has two incorrectly assigned alleles. The large database allows subreads to map with several alleles creating noise in the forensic profile, leading to drop-ins and drop-outs when alleles are being called without prior knowledge. Homozygous loci, like D7S820, D13S317, D16S539 and vWA, are less prone to this issue and show no drop-ins or drop-outs in our dataset. A reason for the better signal-to-noise ratio of homozygous alleles can be found in the fact only one allele has to be detected. The correctly mapped reads at a homozygous allele will give twice the signal compared to the reads mapped at the two alleles form a heterozygous sample. Hence, homozygous samples will more easily rise above the noise allowing better detection.

To study sequencing errors, which can be both random and systematic, consensus sequences were generated using the reads that mapped uniquely at the true alleles (data not shown). With enough coverage, random errors are corrected in a consensus sequence, but systematic errors are not. A frequently observed systematic error found in our data, was a partial deletion in homopolymeric stretches. Several other members of the ONT community have also reported such bias. The extraction of single-base identities inside homopolymeric regions based on the ionic current state proves to be challenging for the base calling algorithm, as the individual bases are hard to discern. Sequencing of such homopolymers has also proven to be difficult for other sequencing technologies (e.g. 454 pyrosequencing and Ion Torrent) [^17–19^]. A possibility to circumvent this STR related homopolymeric sequencing bias, while still generating forensically viable profiles, would be to sequence SNPs (Single Nucleotide Polymorphisms). SNPs based human identification, in contrast to STRs, relies on the detection of only one alternative nucleotide, which should make them less prone the homopolymeric sequencing bias. To evaluate the possibility to detect SNPs, the AmelX subreads were forced to align with the AmelY reference. Besides the obvious six bp deletion, the seven SNVs, which are known to be different between the AmelX and AmelY locus, can be identified. This illustrates the capability of the MinION to identify SNVs and SNPs. Although SNVs/SNPs are not yet routinely used in forensics, interest in forensic use of SNVs has increased over the past several years [^20–22^].

### 4.2. Length-based profiling

The second approach tested in this paper is based on the length of the generated sub-reads, which is an approach very similar to the traditional CE profile generation. The rationale behind this concept is that although sequencing errors might occur, the sequence length might still be correct. A length histogram of the generated sequences was created for each locus. Because of the wide length distribution of the generated subreads, a large number of alleles is included under each histogram. In some cases, the entire range of the frequently observed alleles is covered by the histogram. To determine the mean length and the related SD, a Gaussian curve fitting was performed on these histograms. The only locus where the data allowed a dual fitting was the TH01 locus, although multiple loci were heterozygous. It is clear that the MinION sequencer is prone to generate a high amount of indel errors, which results in a wide length distribution of the subsequent subreads.

### 4.3. Evidential value of the profile

Although the obtained profiles cannot be used in current practice, as they are subpar to the current CE and Illumina sequencing profiles, the RMNE value was calculated. These RMNE values allow us to describe the evidential value of the obtained profiles as such. The RMNE approach calculates the evidential value of a profile without reference to a suspect’s profile or a defense hypothesis. The number of contributors does not need to be estimated. It also allows accounting for drop-outs without the need to estimate the probability that drop-outs have occurred in the profile. As this is no validation study, no knowledge exists that could enable us to estimate the drop-out and drop-in probability. Both the sequence-and the length-based profile have some evidential value, albeit limited. The sequence-based profile contains seven correctly called loci, but also 6 drop-outs and 4 drop-ins, resulting in an RMNE value of 5.32E-07. The evidential value of the length-based profile is much worse (2.6E-4) because the observed lengths encompass too many alleles. The same sample, analyzed with CE, results in a completely correct profile with an RMNE value that is 10E+07 and 10E+10 times smaller compared to the sequence-and length-based profile respectively.

## 5 Conclusion

In this proof-of-principle study the applicability of ONT’s MinION nanopore sequencers for forensic STR profiling was explored. Two approaches were investigated, one based on the sequence length to generate a CE-like profile and the other based on the actual sequence. Both approaches failed to produce a conclusive forensic profile due to a high amount of sequencing errors. Despite this high sequencing error rate, we were able to extract partial forensic STR profiles with some evidential value from the nanopore data. The technique is however still subpar compared with other sequencing techniques and capillary electrophoresis to produce STR profiles. The potential of nanopore sequencing to generate forensic SNP profiles was confirmed as the seven known SNVs that distinguish the X and Y amelogenin from each other could be readily detected.

## 6 Funding

This research was mainly funded by a PhD grant from the Institute for the Promotion of Innovation through Science and Technology in Flanders (IWT-Vlaanderen), awarded to Senne Cornelis (Grant nr°141456) and a PhD grant from the Special Research Fund (BOF) from the Ghent University awarded to Olivier Tytgat (Grant Nr°:BOF18/DOC/200) and Jana Weymaere (Grant Nr°:BOF17/DOC/265)

## 7 Additional information

The authors declare no conflict of interests.

## Acknowledgements

We would like to thank Sarah De Keulenaer, Ellen De Meester from NXTGNT Belgium, Hendrik Vandevoorde and Johan Vandersmissen for their invaluable practical expertise and assistance in the experiments of this study.

